# Variational Autoencoder: An Unsupervised Model for Modeling and Decoding fMRI Activity in Visual Cortex

**DOI:** 10.1101/214247

**Authors:** Kuan Han, Haiguang Wen, Junxing Shi, Kun-Han Lu, Yizhen Zhang, Zhongming Liu

## Abstract

Goal-driven and feedforward-only convolutional neural networks (CNN) have been shown to be able to predict and decode cortical responses to natural images or videos. Here, we explored an alternative deep neural network, variational auto-encoder (VAE), as a computational model of the visual cortex. We trained a VAE with a five-layer encoder and a five-layer decoder to learn visual representations from a diverse set of unlabeled images. Inspired by the “free-energy” principle in neuroscience, we modeled the brain’s bottom-up and top-down pathways using the VAE’s encoder and decoder, respectively. Following such conceptual relationships, we used VAE to predict or decode cortical activity observed with functional magnetic resonance imaging (fMRI) from three human subjects passively watching natural videos. Compared to CNN, VAE resulted in relatively lower accuracies for predicting the fMRI responses to the video stimuli, especially for higher-order ventral visual areas. However, VAE offered a more convenient strategy for decoding the fMRI activity to reconstruct the video input, by first converting the fMRI activity to the VAE’s latent variables, and then converting the latent variables to the reconstructed video frames through the VAE’s decoder. This strategy was more advantageous than alternative decoding methods, e.g. partial least square regression, by reconstructing both the spatial structure and color of the visual input. Findings from this study support the notion that the brain, at least in part, bears a generative model of the visual world.

## Introduction

Humans readily make sense of the visual surroundings through complex neuronal circuits. Understanding the human visual system requires not only measurements of brain activity but also computational models built upon hypotheses about neural computation and learning (Kietzmann et al., 2017). Models that truly reflect the brain’s mechanism of natural vision should be able to explain and predict brain activity given any visual input (encoding), and decode brain activity to infer visual input (decoding) (Naselaris et al., 2011). In this sense, evaluating the models’ encoding and decoding performance serves to test and compare hypotheses about how the brain learns and organizes visual representations (Wu et al., 2006).

In one class of hypotheses, the visual system consists of feature detectors that progressively extract and integrate features for pattern recognition. For example, Gabor and wavelet filters model low-level features (Hubel and Wiesel, 1962; van Hateren and van der Schaaf, 1998), and explain responses in early visual areas (Kay et al., 2008; Nishimoto et al., 2011). In contrast, convolutional neural networks (CNNs) encode hierarchical features in a feedforward model (LeCun et al., 2015), and support high performance in image recognition (He et al., 2016; Krizhevsky et al., 2012; Simonyan and Zisserman, 2014). Recent studies have shown that CNNs bear similar representations as does the brain (Cichy et al., 2016; Khaligh-Razavi and Kriegeskorte, 2014), and yield high performance in neural encoding and decoding of natural vision (Eickenberg et al., 2017; Guclu and van Gerven, 2015; Horikawa and Kamitani, 2017; Wen et al., 2017b; Yamins et al., 2014). For these reasons, supervised CNN models are gaining attention as favorable models of the visual cortex (Kriegeskorte, 2015; Yamins and DiCarlo, 2016). However, biological learning is not always supervised towards a single goal, but often unsupervised (Barlow, 1989). The visual cortex has not only feedforward but also feedback pathways (Bastos et al., 2012; Salin and Bullier, 1995). As such, CNNs are not an ideal computational account of human vision.

In another class of hypotheses, the brain builds a causal model of the world, through which it tries to infer what generates the sensory input in order to make proper perceptual and behavioral decisions (Friston, 2010). In this scenario, the brain behaves as an inference machine: recognizing and predicting visual input through “analysis by synthesis” (Yuille and Kersten, 2006). The brain’s bottom-up process infers the “cause” of the input, and its top-down process predicts the input (Fig. 1.A). Both processes are optimized by learning from visual experience in order to avoid the “surprise” or error of prediction (Friston and Kiebel, 2009; Rao and Ballard, 1999). This hypothesis takes into account both feedforward and feedback pathways. It aligns with the humans’ ability to construct mental images, and offers a basis for unsupervised learning. Thus, it is compelling for both computational neuroscience (Bastos et al., 2012; Friston, 2010; Rao and Ballard, 1999; Yuille and Kersten, 2006) and artificial intelligence (Hinton et al., 1995; Lotter et al., 2016; Mirza et al., 2016).

**Figure 1.**
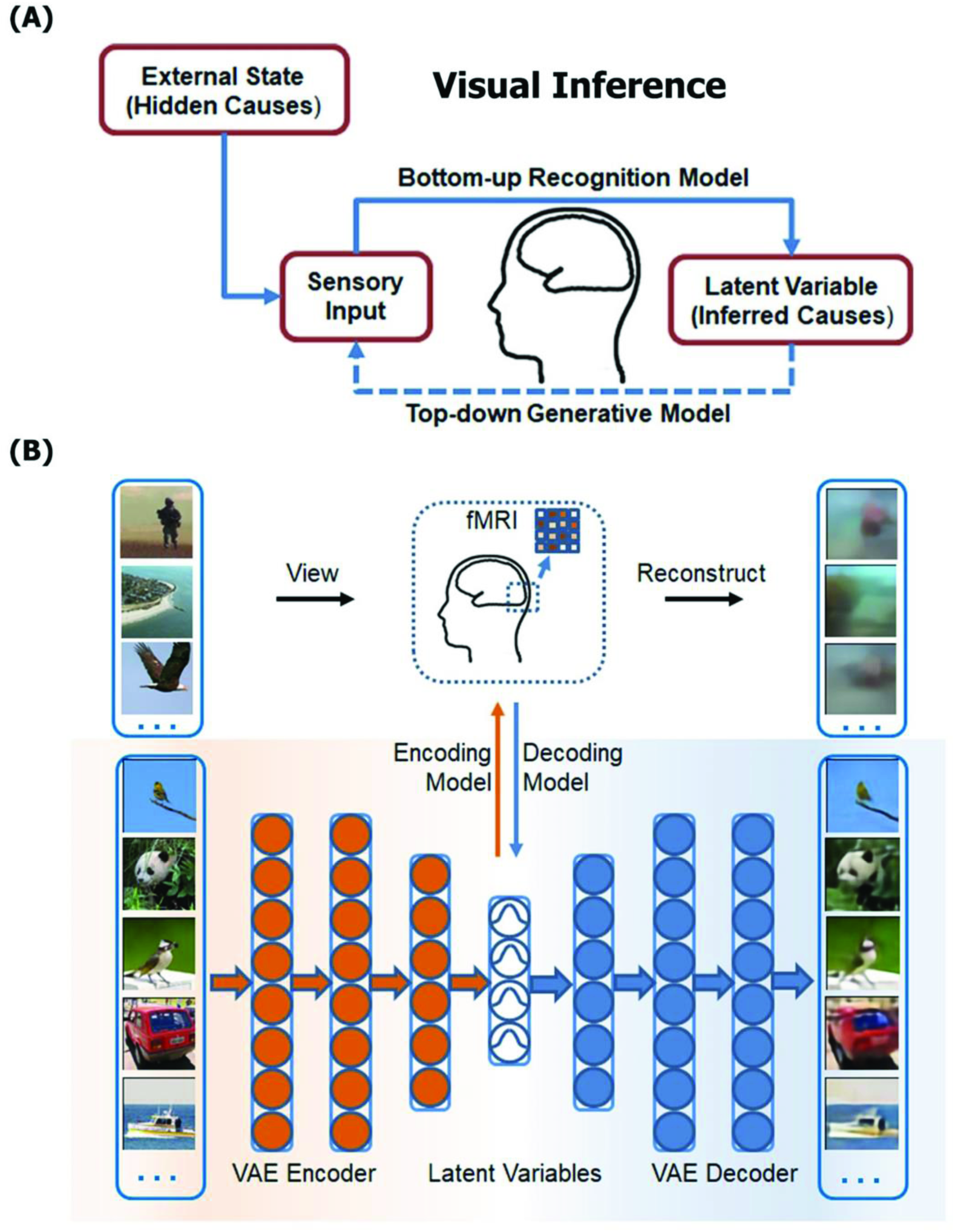
A Bayesian model of human vision. **(A) The brain models the causal structure of the world for visual inference**. The brain analyzes the sensory input to infer the hidden causes of the input through its bottom-up processes, and predicts the sensory input through its top-down processes. **(B) Encoding and decoding visually-evoked cortical fMRI responses by using VAE as a model of the visual cortex**. For encoding, cortical responses to any visual stimuli are predicted as a linear projection of the VAE responses to the same stimuli. For decoding, visual stimuli were reconstructed by first estimating the VAE’s latent variables as a linear function of the fMRI responses, and then generating pixel patterns from the estimated through the VAE’s decoder.

In line with this notion, variational auto-encoder (VAE) uses independent “latent” variables to code the causes of the visual world (Kingma and Welling, 2013). VAE learns the latent variables from images via an encoder, and samples the latent variables to generate new images via a decoder. Both the encoder and the decoder are neural networks trainable without supervision from unlabeled images (Doersch, 2016). Hypothetically, VAE offers a potential model of the brain’s visual system, and may enable an effective way to decode brain activity during either visual perception or imagery (Du et al., 2017; Güçlütürk et al., 2017; Shen et al., 2017; van Gerven et al., 2010a). In this study, we explored the relationships between the VAE’s learning objective and the “free-energy” principle in neuroscience (Friston, 2010). To test VAE as a model of the visual cortex, we built and trained a VAE for unsupervised learning of visual representations, and evaluated the use of VAE for encoding and decoding functional magnetic resonance imaging (fMRI) responses to naturalistic movie stimuli (Fig. 1B).

## Methods and Materials

### Theory: variational auto-encoder

In general, VAE is a type of deep neural networks that learns representations from complex data without supervision (Kingma and Welling, 2013). A VAE includes an encoder and a decoder, both of which are neural nets. The encoder learns latent variables from the input, and the decoder generates outputs similar to the input from samples of the latent variables. Given large training datasets, the encoder and the decoder are trained altogether by minimizing the reconstruction loss and the Kullback-Leibler (KL) divergence between the distributions of the latent variables and independent standard normal distributions (Doersch, 2016). When the input data are natural images, the latent variables represent the hidden causes or attributes of the images.

Mathematically, let ***z*** be the latent variables and ***x*** be an image. The encoder parameterized with ***φ*** infers ***z*** from ***x***, and the decoder parameterized with ***θ*** generates ***x*** from ***z***. In VAE, both ***z*** and ***x*** are random variables. The likelihood of ***x*** given ***z*** under the generative model ***θ*** is denoted as *p*_***θ***_(***x***|***z***). The probability of ***z*** given ***x*** under the inference model ***φ*** is denoted as *q*_***φ***_(***z***|***x***). The marginal likelihood of data can be written as the following form.

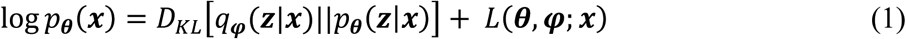

Since the KL divergence in Equation (1) is non-negative, *L*(***θ, φ; x***) can be regarded as the lower-bound of data likelihood and also be rewritten as Eq. (2). For VAE, the learning rule is to optimize ***θ*** and ***φ*** by maximizing *L*(***θ, φ; x***) given the training samples of ***x***.

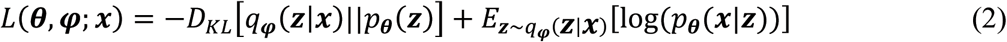

In this objective function, the first term is the KL divergence between the distribution of ***z*** inferred from ***x*** and the prior distribution of ***z***, both of which are assumed to follow a multivariate normal distribution. The second term is the expectation of the log-likelihood that the input image can be generated by the sampled values of ***z*** from the inferred distribution *q*_***φ***_(***z***|***x***). When *q*_***φ***_(***z***|***x***) is a multivariate normal distribution with unknown expectations ***μ*** and variances ***σ***^2^, the objective function is differentiable with respect to (***θ, φ, μ, σ***) with the re-parameterization trick (Kingma and Welling, 2013). The parameters in VAE could be optimized iteratively using gradient-descent algorithms with the Adam optimizer (Kingma and Ba, 2014).

Similar concepts have been put forth in computational neuroscience theories, for example the free-energy principle (Friston, 2010). In the free-energy principle, the brain’s perceptual system includes bottom-up and top-down pathways. Like the encoder in VAE, the bottom-up pathway infers the causes of sensation as probabilistic representations. Like the decoder in VAE, the top-down pathway predicts the sensation from its causes inferred by the brain. Both the bottom-up and top-down pathways are shaped by experiences, such that the brain infers the causes of the sensory input, and generates the sensory prediction that matches the input with the minimal error or surprise. Mathematically, the learning objectives in both VAE and the free energy principle are similar, both aiming to minimize the lower bound of the marginal likelihood (or the free energy), which depends on the difference (or the KL divergence) between the inferred and hidden causes of sensory data while maximizing the likelihood of the sensory data given the inferred causes.

### Training VAE with diverse natural images

We designed a VAE with 1,024 latent variables, and the encoder and the decoder were both convolutional neural nets with five hidden layers (Fig. 2A). Each convolutional layer included nonlinear units with a Rectified Linear Unit (ReLU) function (Nair and Hinton, 2010), except the last layer in the decoder where a sigmoid function was used to generate normalized pixel values between 0 and 1. The model was trained on the ImageNet ILSVRC2012 dataset (Russakovsky et al., 2015). Training images were resized to 128 × 128 × 3. To enlarge the amount of training data, the original training images were randomly flipped in the horizontal direction, resulting in >2 million training samples in total. The training data were divided into mini-batches with a batch size of 200. For each training example, the pixel intensities were normalized to [0, 1]; the normalized intensity was viewed as the probability of color emission (Gregor et al., 2015). To train the VAE, the Adam optimizer (Kingma and Ba, 2014) was used with a learning rate of 1e-4, as implemented *in PyTorch* (http://pytorch.org/).

**Figure 2.**
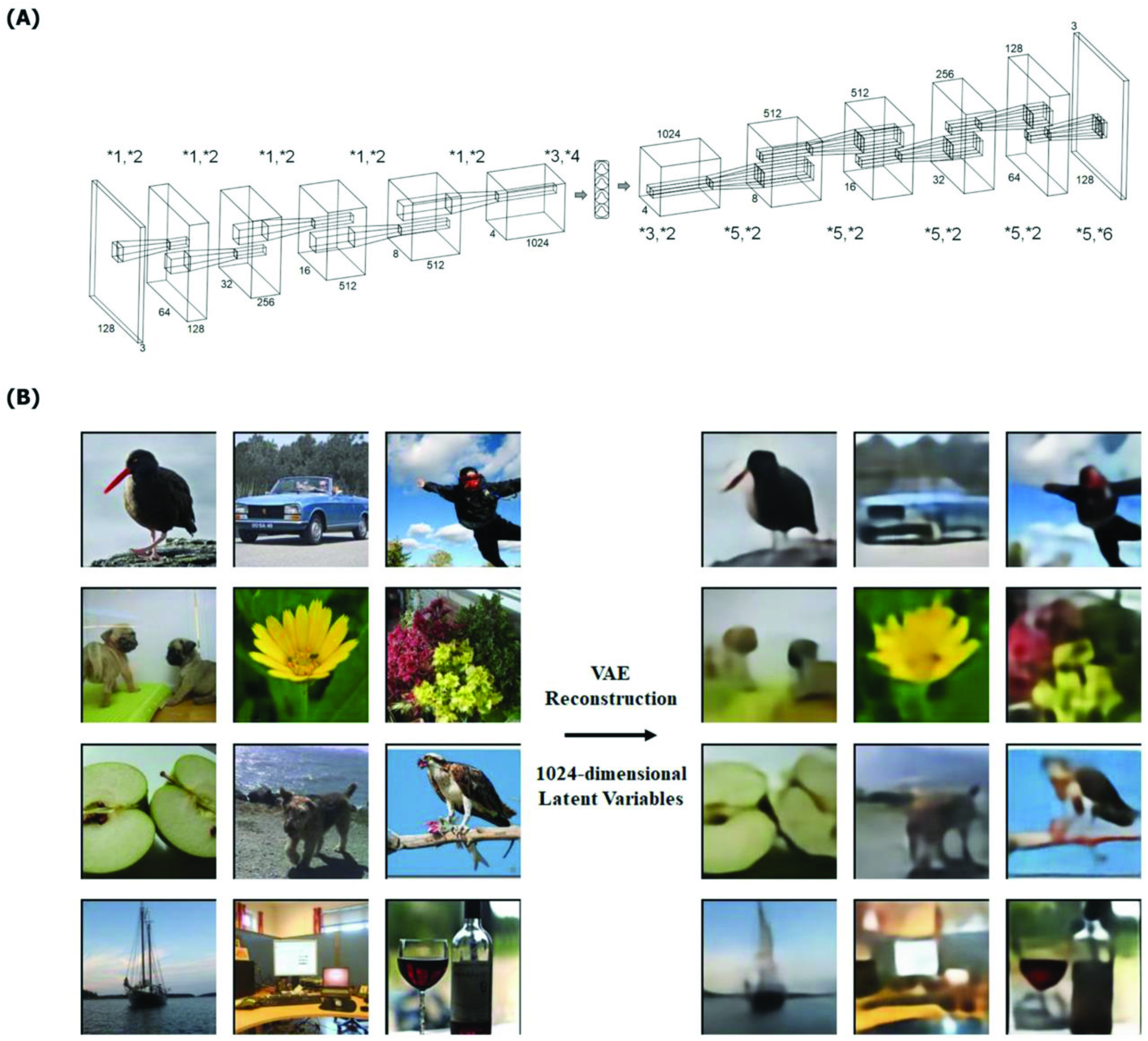
Variational auto-encoder. **(A) The architecture of VAE for natural images**. The encoder and the decoder both contained 5 layers. The dimension of latent variables was 1024. Operations were defined as: *1 convolution (kernel size=4, stride=2, padding=1), *2 rectified nonlinearity, *3 fully connected layer, *4 re-parameterization, *5 transposed convolution (kernel size = 4, stride = 2, padding = 1), *6 sigmoid nonlinearity. **(B) Reconstruction of natural images by VAE**.>For any image (left), its information was encoded by 1024 latent variables by passing it through the VAE encoder. From the latent variables, the VAE decoder generated an image (right) as the reconstruction of the input image, despite blurred details.

### Experimental data

Three healthy volunteers (all female, age: 23–26) participated in this study with informed written consent according to a research protocol approved by the Institutional Review Board at Purdue University. All experiments were performed according to the guidelines and regulations in the protocol. As described in detail elsewhere (Wen et al., 2017b), the experimental design and data were briefly summarized as below. Each subject watched a diverse set of natural videos for a total length up to 13.7 hours. The videos were downloaded from Videoblocks and YouTube, and then were separated into two independent sets. One data set was for training the models to predict the fMRI responses based on the input video (i.e. the encoding models) or the models to reconstruct the input video based on the measured fMRI responses (i.e. the decoding models). The other data set was for testing the trained encoding or decoding models. The videos in the training and testing datasets were independent for unbiased model evaluation. Both the training and testing movies were further split into 8-min segments. Each segment was used as visual stimulation (20.3^o^ × 20.3^o^) along with a central fixation cross (0.8^o^ × 0.8^o^) presented via an MRI-compatible binocular goggle during a single fMRI session. The training movie included 98 segments (13.1 hours) for Subject 1, and 18 segments (1.6 hours) for Subject 2 & 3. The testing movie included 5 segments (40 mins in total). Each subject watched the testing movie 10 times. All five segments of the testing movie were used to test the encoding model. One of the five segments of the testing movie was used to test the decoding models for visual reconstruction, because this segment contained video clips that were continuous over relatively long periods (mean±std: 13.3±4.8 s).

MRI/fMRI data were collected from a 3-T MRI system, including anatomical MRI (T_1_ and T_2_ weighted) of 1mm isotropic spatial resolution, and blood oxygenation level dependent (BOLD) fMRI with 2-s temporal resolution and 3.5mm isotropic spatial resolution. The fMRI data were registered onto anatomical MRI data, and were further co-registered on a cortical surface template (Glasser et al., 2013). The fMRI data were preprocessed with the minimal preprocessing pipeline released for the human connectome project (Glasser et al., 2013).

### VAE-based encoding models

After training, VAE extracted the latent representation of any video by a feed-forward pass of every video frame into the encoder, and reconstructed every video frame by a feedback pass of the latent representation into the decoder. To predict cortical fMRI responses to the video stimuli, an encoding model was defined separately for each voxel as a linear regression model. The voxel-wise fMRI signal was estimated as a linear combination of all unit activities in both the encoder and the decoder given the input video. Every unit activity in VAE was convolved with a canonical hemodynamic response function (HRF). For dimension reduction, PCA was applied to the HRF-convolved unit activity for each layer, keeping 99% of the variance of the layer-wise activity given the training movies. Then, the layer-wise activity was concatenated across layers; PCA was applied again to the concatenated activity to keep 99% of the variance of the activity from all layers given the training movies. See details in our earlier paper (Wen et al., 2017a). Following the dimension reduction, the principal components of unit-activity were down-sampled by the sampling rate of fMRI and were used as the regressors to predict the fMRI signal at each voxel through a linear regression model specifically estimated for the voxel.

The voxel-wise regression model was trained with the fMRI data during the training movie. Mathematically, for any training sample, let ***x***^*(j)*^ be the visual input at the *j*-th time point, *y*_***i***_^*(j)*^ be the fMRI response at the i-th voxel, ***z***^*(j)*^ be a vector representing the predictors for the fMRI signal derived from ***x***^*(j)*^ through VAE, as described above. The voxel-wise regression model is expressed as Eq. (4).

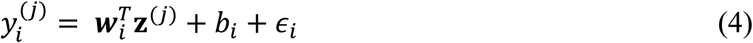

where ***w***_*i*_ is a column vector representing the regression coefficients, ***b***_*i*_ is the bias term, and ***∊***_*i*_ is the error unexplained by the model. The linear regression coefficients were estimated using least-squares estimation with L_2_-norm regularization, or minimizing the loss function as Eq. (5).

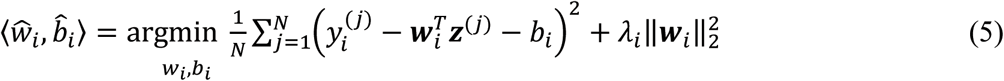

where *N* is the number of training samples. The regularization parameter *λ*_*i*_ was optimized for each voxel to minimize the loss in three-fold cross-validation. Once the parameter *λ*_*i*_ was optimized, the model was refitted with the entire training dataset and the optimized parameter.

### Evaluation of encoding performance

After the above model training, we tested the voxel-wise encoding models with the testing movies, which were different from the training movies to ensure unbiased model evaluation. For each voxel, the encoding performance was evaluated as the correlation between the measured and predicted fMRI responses to the testing movie. The significance of the correlation was assessed by using a block-wise permutation test with a block size of 24-sec and 100,000 permutations and corrected at false discovery rate (FDR) *q* < 0.01, as described in our earlier papers (Shi et al., 2017; Wen et al., 2017b).

We compared the encoding performance against the so-called “noise ceiling”. It indicated the upper-limit of predictability given the presence of “noise” unrelated to the visual stimuli (David and Gallant, 2005; Nili et al., 2014). The noise ceiling was estimated using the method described elsewhere (Kay et al., 2013). Briefly, the noise was assumed to follow a Gaussian distribution with a zero mean and an unknown variance that varied across voxels. The response and the noise were assumed to be independent and additive. The variance of the noise was estimated as the squared standard error of the mean of fMRI signal (averaged across the 10 repeated sessions of each testing movie). The variance of the response was taken as the difference between the variance of the data and the variance of the noise. From the signal and noise distributions, the samples of the response and the noise were drawn by Monte Carlo simulation for 1,000 random trials. For each trial, the signal was simulated as the sum of the simulated response and noise; its correlation coefficient with the simulated response was calculated. This resulted in an empirical distribution of the correlation coefficient for each voxel. We calculated the mean of the distribution as the noise ceiling *r*_*NC*_, and interpreted it as the upper limit of the prediction accuracy for any encoding model.

In terms of the encoding performance, we compared the encoding models based on VAE against those based on CNN, which have been explored in recent studies (Eickenberg et al., 2017; Guclu and van Gerven, 2015; Wen et al., 2017b). For this purpose, we used a 18-layer residual network (ResNet-18) (He et al., 2016). Relative to AlexNet (Krizhevsky et al., 2012) or ResNet-50 (He et al., 2016), ResNet-18 had an intermediate level of architectural complexity in terms of the number of layers and the total number of units. Thus, ResNet-18 was a suitable benchmark for comparison with VAE, which had a comparable level of complexity. Briefly, ResNet-18 consisted of 18 hidden layers organized into 6 blocks. The 1^st^ block was a convolutional layer followed by max-pooling; the 2^nd^ through 5^th^ blocks were residual blocks, each being a stack of convolutional layers with a shortcut connection; the 6^th^ block performed the multinomial logistic regression for classification. Typical to CNNs, ResNet-18 encoded increasingly complex visual features from lower to higher layers.

We similarly built and trained voxel-wise regression models to project the representations in ResNet-18 to voxel responses in the brain, using the same training procedure and data as above for VAE-based encoding models. Then, we compared VAE against CNN (ResNet-18) in terms of the encoding performance evaluated in the level of voxels or regions of interest (ROI). In the voxel level, the encoding accuracy was converted from the correlation coefficient to the z score by the Fisher’s z-transform for the VAE or CNN-based encoding models. Their difference in the voxel-wise z score was calculated by subtraction. For the ROI-level comparison, multiple ROIs were selected from existing cortical parcellation (Glasser et al., 2016), including V1, V2, V3, V4, lateral occipital (LO), middle temporal (MT), fusiform face area (FFA), para-hippocampal place area (PPA) and temporoparietal junction (TPJ). In each ROI, the correlation coefficient of each voxel was divided by the noise ceiling, and then averaged over all voxels in the ROI. The average prediction accuracy of each ROI was compared between VAE and ResNet-18.

### VAE-based decoding of fMRI for visual reconstruction

We trained and tested the decoding model for reconstructing visual input from distributed fMRI responses. The model contained two steps: 1) transforming the fMRI response pattern to the latent variables in VAE through a linear regression model, and 2) transforming the latent variables to pixel patterns through the VAE’s decoder. Here we used a cortical mask that covered the visual cortex, and used the voxels within the mask as the input to the decoding model as in our previous study (Wen et al., 2017b).

Let ***y***^*(j)*^) be a column vector representing the measured fMRI map at the *j*-th time point, and ***z***^*(j)*^ be a column vector representing the means of the latent variables given the visual input ***x***^*(j)*^. As Eq. (6), a multivariate linear regression model was defined to predict ***z***^*(j)*^ given ***y***^*(j)*^.

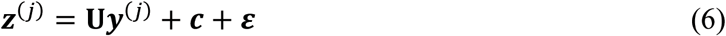

where **U** is a weighting matrix representing the regression coefficients to transform the fMRI map to the latent variables, ***c*** is the bias term, and ***ε*** is the error term unexplained by the model. This model was estimated based on the data during the training movie.

To estimate parameters of the decoding model, we minimized the objective function as Eq. (7) with L1-regularized least-squares estimation to prevent over-fitting.

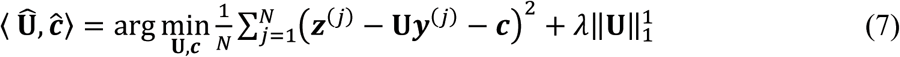

where *N* is the number of data samples used for training the model. The regularization parameter, *λ*, was optimized to minimize the loss in three-fold cross-validation. To solve Eq. (7), we used the mini-batch stochastic gradient-descent algorithm with a batch size of 100 and a learning rate of 1e-7.

It follows that the testing movie was reconstructed frame by frame by passing the estimated latent variables through the decoder in VAE, as expressed by Eq. (8)

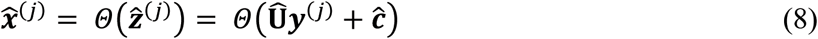

where *Θ* is the nonlinear mapping from latent variables to the visual reconstruction defined by the VAE decoder.

### Evaluation of decoding performance

To evaluate the decoding performance in visual reconstruction, we calculated the Structural Similarity index (SSIM) (Wang et al., 2004) between every reconstructed video frame and the true video frame, yielding a measure of the similarity in the pattern of pixel intensity. The SSIM was further averaged across all video frames in the testing movie.

In addition, we evaluated how well movie reconstruction preserved the color information in the original movie. For this purpose, the color information at each pixel was converted from the RGB values to a single hue value. The hue maps of the reconstructed movie frames were compared with those of the original movie frames. Their similarity was evaluated in terms of the circular correlation (Berens, 2009; Jammalamadaka and Sengupta, 2001). Hue values in the same frame were represented as a vector and the hue vectors were concatenated sequentially across all video frames to represent the color information in the entire movie. The circular correlation in the concatenated hue vector between the original and reconstructed frames was calculated for each subject. The statistical significance of the circular correlation between the reconstructed and original color was tested by using the block-permutation test with 24-sec block size and 100,000 times permutation (Adolf et al., 2014).

We also compared the performance of the VAE-based decoding method with a previously published decoding method (Cowen et al., 2014). In this alternative method (Cowen et al., 2014), we applied PCA to the training movie and obtained its principal components (or eigen-images), which explained 99% of the variance in the pixel pattern of the movie. The partial least square regression (PLSR) (Tenenhaus et al., 2005) was used to estimate the linear transformation from fMRI maps to eigen-images given the training movie. Using the estimated PLSR model, the fMRI data during the testing movie was converted to representations in eigen-images, which in turn were recombined to reconstruct the visual stimuli (Cowen et al., 2014). As a variation of this PLSR-based model, we also explored the use of L1-norm regularized optimization to estimate the linear transform from fMRI maps to eigen-images, in a similar way as used for the training of our decoding model (see Eq. (7)). As such, the training procedure was identical for both methods, except that the feature space for decoding was different: the latent variables for VAE, and eigen-images for PLSR.

Moreover, we also explored whether the VAE-based decoding models could be generalized across subjects. For this purpose, we used the decoding model trained from one subject to decode the fMRI data observed from the other subjects while watching the testing movie.

## Results

### VAE provided vector representations of natural images

By design, VAE aimed to form a compressed and generalized vector representation of any natural image. In VAE, the encoder converted any natural image into 1,024 independent latent variables; the decoder reconstructed the image from the latent variables (Fig. 2.A). After training it with >2 million natural images in a wide range of categories, the VAE could regenerate natural images without a significant loss in image content, structure and color, albeit blurred details (Fig. 2B). The VAE-generated images showed comparable quality for different types of input images (Fig. 2.B). As such, the latent representations in VAE were generalizable across various types of visual objects, or their combinations.

### VAE predicted movie-induced cortical responses

Given natural movies as visual input, we further asked to what extent the model dynamics in VAE could be used to model and predict the movie-induced cortical responses. Specifically, a linear regression model was trained separately for each voxel by fitting the voxel response to a training movie as a linear combination of the VAE’s unit responses to the same movie. Then, the trained voxel-wise encoding model was tested with a new testing movie (not used for training) to evaluate the model’s prediction accuracy (i.e. the correlation between the predicted and measured fMRI responses). For a large area in the visual cortex (Fig. 3), the VAE-based encoding models could predict the movie-evoked responses with statistically significant accuracy (FDR q<0.01). In particular, early visual areas (V1/V2/V3) showed the highest prediction accuracy, whereas the prediction accuracy was relatively lower for higher visual areas along the ventral or dorsal stream (Fig. 3). The VAE-predictable areas were relatively larger when more data (~12-hour movie) were used for training the encoding models in Subject 1 than Subject 2 & 3 for whom less training data (2.5-hour movie) were available.

**Figure 3.**
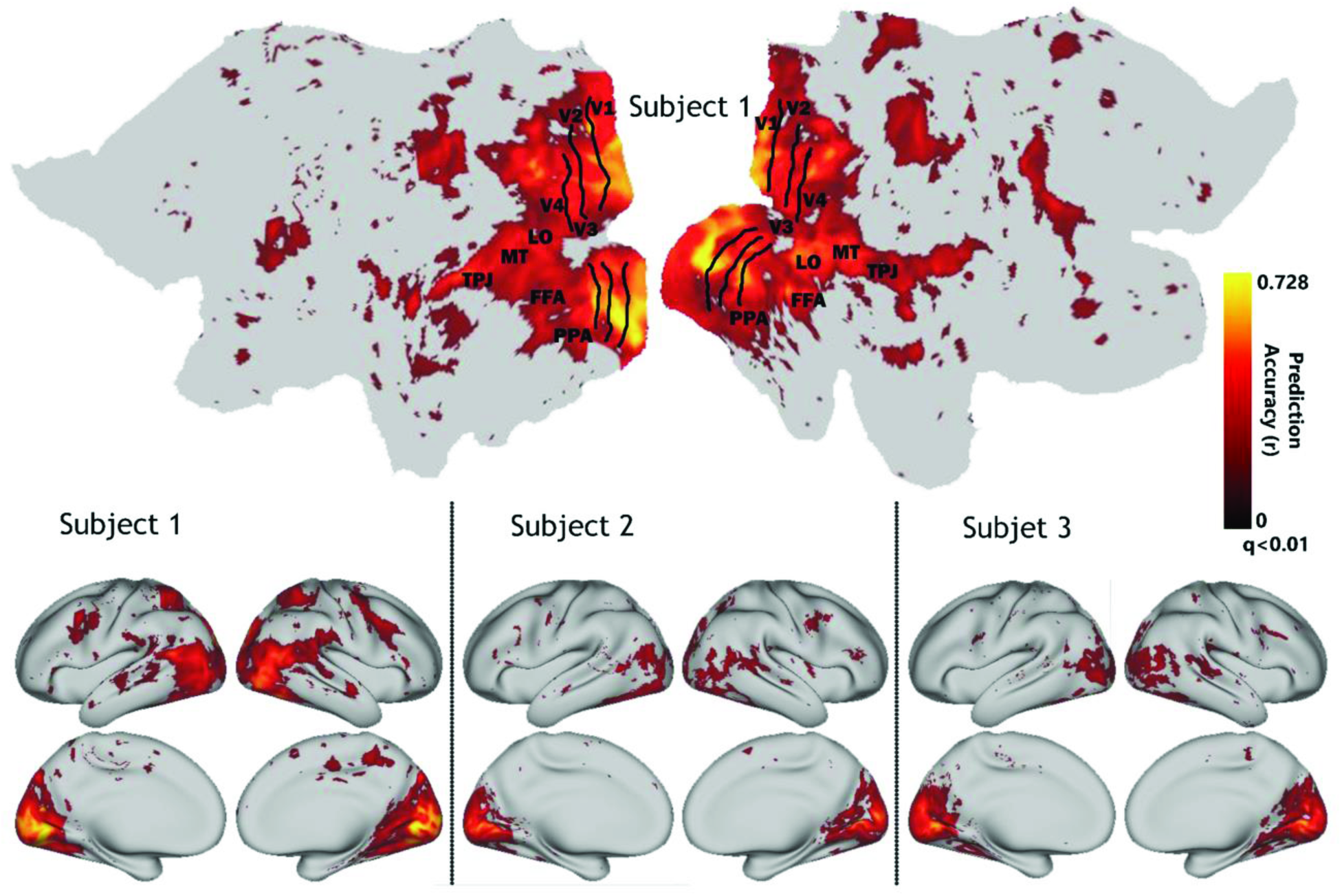
Prediction accuracy with VAE-based encoding models. The accuracy was measured by the Pearson’s correlation coefficient (r) between the model-predicted response and the actual fMRI response. The map shows the r value averaged across the five testing movies. The map was thresholded by statistical significance (FDR q<0.01). The results are shown on the flattened (only for Subject 1) and inflated cortical surfaces for every subject.

### Comparing encoding performance for VAE vs. CNN

We further compared VAE against CNN, which was found to predict cortical responses to natural picture or video stimuli (Eickenberg et al., 2017; Guclu and van Gerven, 2015; Wen et al., 2017b). Fig. 4 shows the prediction accuracies at different ROIs, given the encoding models based on VAE and ResNet-18 – a particular type of CNN. For VAE, the encoding performance was the highest in V1, and decreased progressively towards higher visual areas. ResNet outperformed VAE in all ROIs, especially for higher visual areas along the ventral stream (e.g. FFA/PPA/TPJ), but marginally for early visual areas. Similar findings were observable in the voxel level. Fig. 5 shows the voxel-wise prediction accuracy for VAE or ResNet. ResNet outperformed VAE for most of the visual cortex. Their difference (by subtraction) was much more notable in the ventral-stream areas than early visual areas or those in the dorsal-stream areas. In sum, VAE was in general less predictive of visual cortical activity than was CNN.

**Figure 4.**
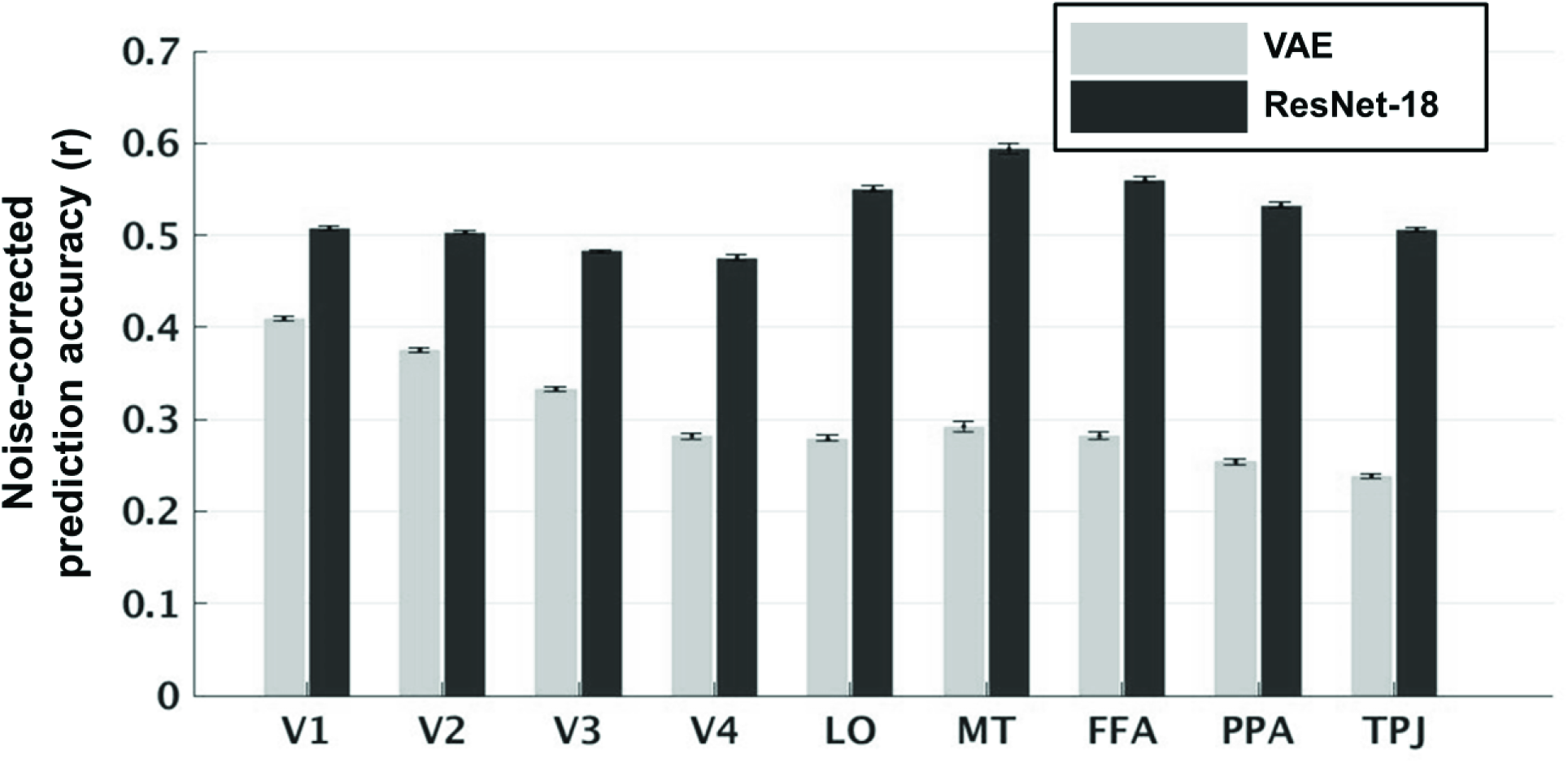
The ROI-level encoding performance of VAE vs. ResNet-18. For either VAE (light array) or ResNet-18 (dark gray), the encoding model’s accuracy in predicting the fMRI response to the testing movie was normalized by the noise ceiling and summarized for each of the nine predefined ROIs. Arranged from the left to the right, individual ROIs are located in increasingly higher levels of the visual hierarchy. The bar chart was based on the mean±SEM (standard error of the mean) of the voxel-wise prediction accuracy (divided by the noise ceiling) averaged across all the voxels in each ROI, and across different testing movies and subjects.

**Figure 5.**
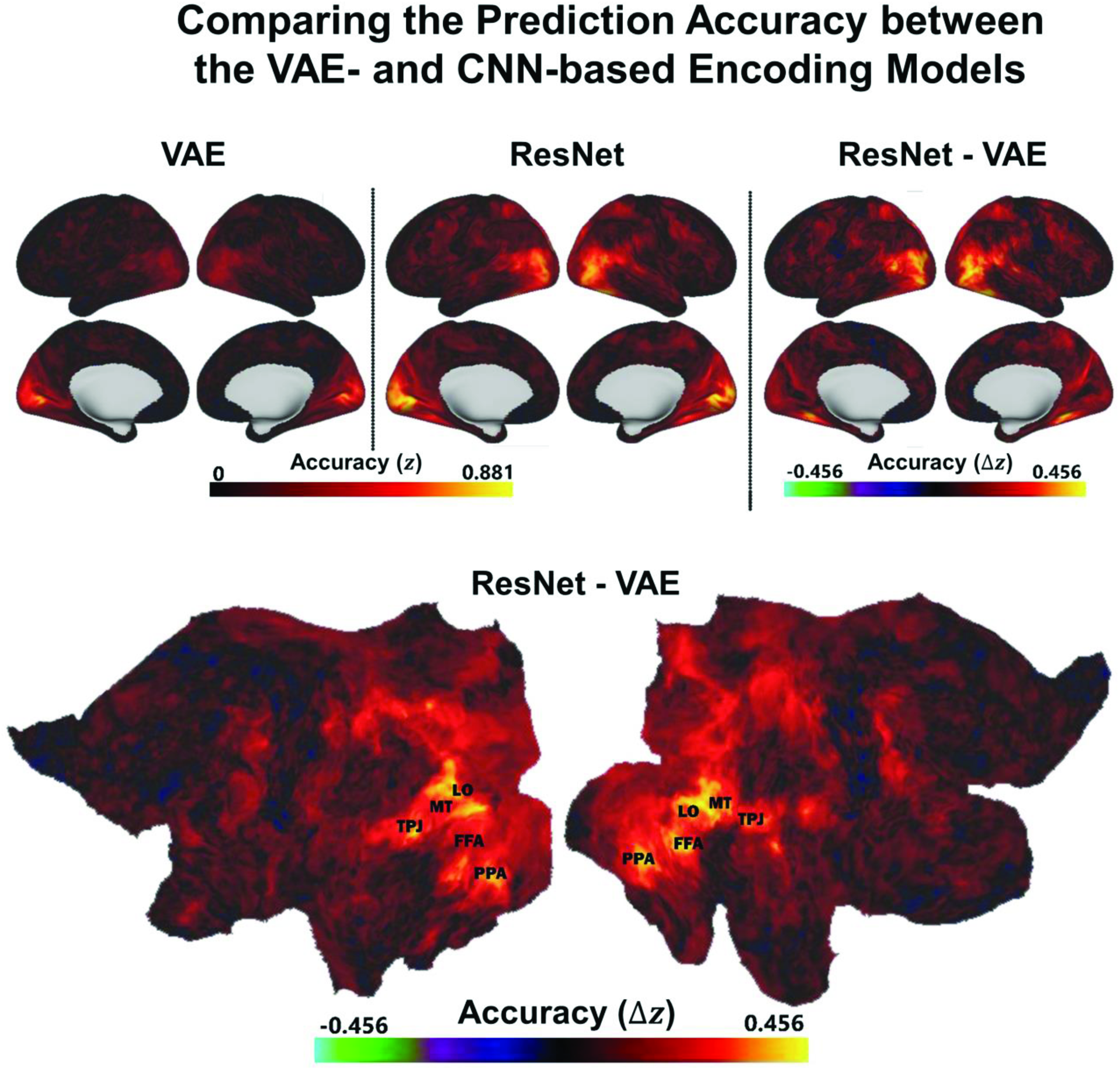
Encoding performance with VAE vs. ResNet-18. The prediction accuracy (the z-transformed correlation between the predicted and measured fMRI responses) is displayed on inflated cortical surfaces for the encoding models based on VAE (top-left) and ResNet-18 (top-middle). Their difference (ResNet – VAE) in the prediction accuracy issplayed on both inflated (top-right) and flattened (bottom) cortex.

### Direct visual reconstruction by decoding fMRI activity

We further explored the use of VAE for decoding the fMRI activity to reconstruct the visual input. For this purpose, a decoding model was trained and used to convert the fMRI activity to the VAE’s latent-variable representation, which was in turn converted to a pixel pattern through the VAE’s decoder. In comparison with the original videos, Fig. 6 shows the visual input reconstructed from fMRI activity based on VAE and the decoding models, which were trained and tested with data from either the same or different subjects. Although the visual reconstruction was too blurry to fully resolve details or discern visual objects, it captured basic information about the dynamic visual scenes, including the coarse position and shape of objects, and the rough color and contrast. The quality of visual reconstruction was better when the decoding models were trained and tested for the same subject than for different subjects.

**Figure 6.**
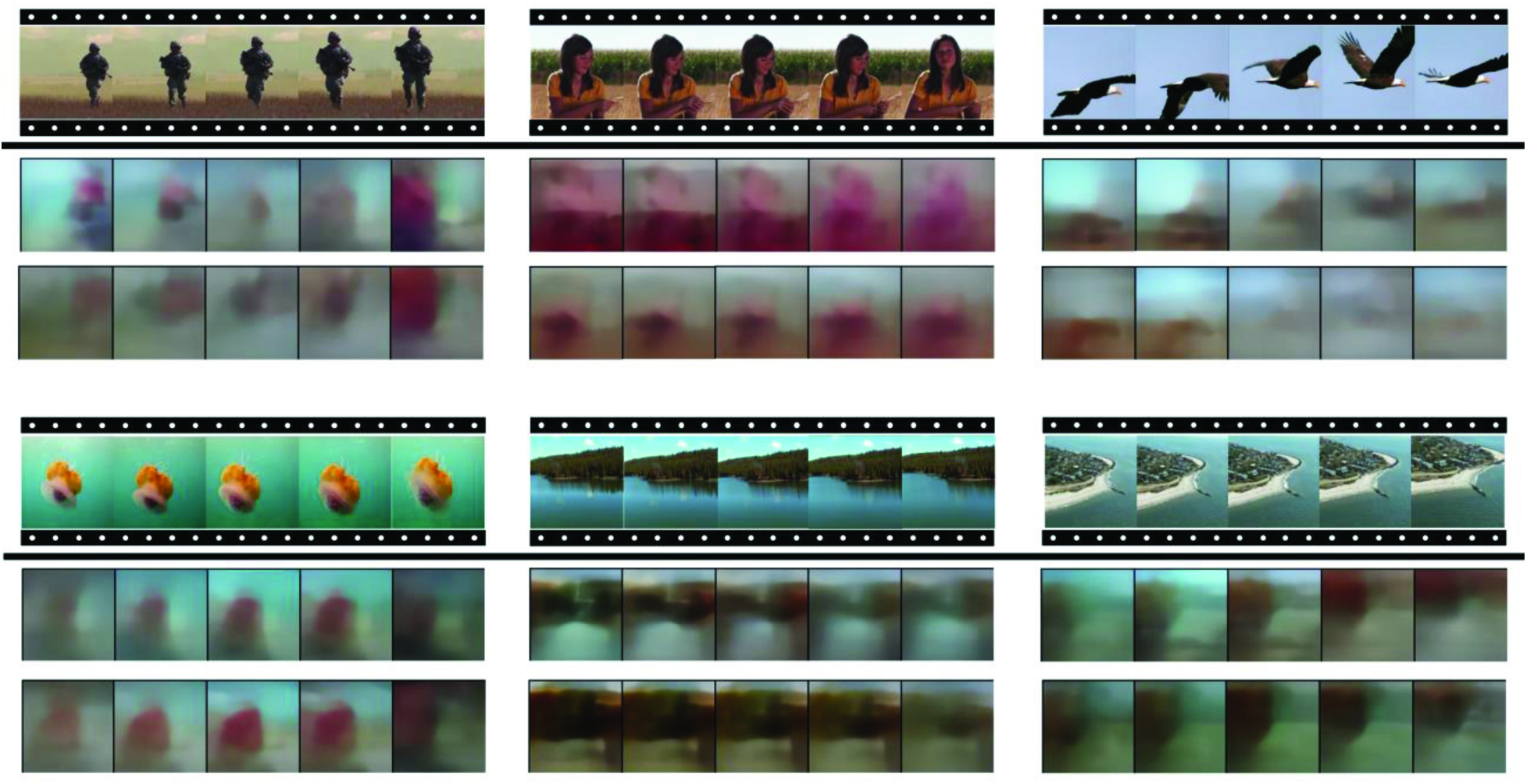
Visual reconstruction based on VAE and fMRI. For each of the 6 example video clips in the testing movie, the top row shows the original video frames, the middle or bottom rows show the frames reconstructed from the measured fMRI responses, based on VAE and the decoding model trained and tested within the same subject (Subject 1), or across different subjects, respectively. The cross-subject decoding models were trained with data from Subject 1, but tested on data from Subject 2 (the top 3 clips) or Subject 3 (the bottom 3 clips).

We assessed the quality of visual reconstruction by quantifying the structural similarity (as SSIM) (Wang et al., 2004) between the reconstructed and original movies. The VAE-based decoding method yielded a much higher SSIM (about 0.5) than the eigen-image-based benchmark models with either partial least squares regression (Cowen et al., 2014) or L1-regularized linear regression (Fig. 7A). Moreover, the VAE-based visual reconstruction preserved the color information and its dynamics in the movie, showing statistically significant (permutation test, p < 0.001) correlations in color index (Hue-value) around 0.25.

**Figure 7.**
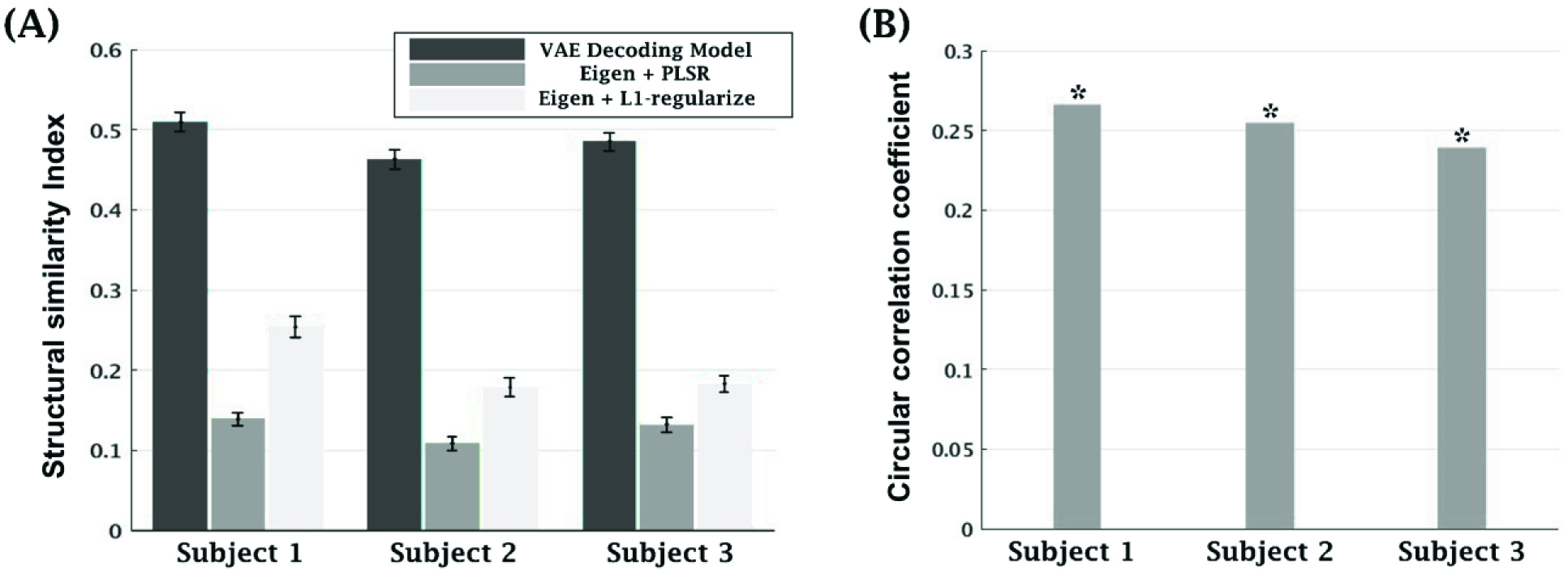
Quantitative evaluation of visual reconstruction. **(A) Comparison of structural similarity index (SSIM)**. SSIM scores of VAE-based decoding model and eigen-image-based models were compared for all 3 subjects. Each bar shows the mean±SE SSIM score over all frames in the testing movie. **(B) Correlation in color (hue-value)**. The (circular) correlation between the original and reconstructed hue components was calculated and evaluated for statistical significance with permutation test (*, p<0.001).

Lastly, we decoded the fMRI activity of Subject 2 or 3 based on the decoding model trained from Subject 1 (Fig. 6). By visual inspection, the quality of reconstruction was lower as the model was trained and tested with different subjects. Nevertheless, the reconstruction was qualitatively similar to what was attained with the decoding models trained/tested with the same subject, while preserving basic patterns in the original video frames (Fig. 6). Therefore, the VAE-based decoding model was transferable across subjects.

## Discussion

We designed and trained a variational autoencoder (VAE) to learn visual representations by reconstructing diverse natural images. Unlike CNN, VAE does not require labeled samples, and can be trained by unsupervised learning based on variational inference (Kingma and Welling, 2013). Non-observable causes of visual input are inferred through joint optimization of bottom-up and top-down components in VAE. In line with the notion of Bayesian brain (Yuille and Kersten, 2006), the bottom-up process infers the causes of any given visual input, and the top-down process generates a “virtual version” of the visual input based on the inferred cause. As a candidate model of the visual cortex, VAE is able to decode fMRI scans for visual reconstruction, and predict fMRI responses to video stimuli. The decoding strategy is generative in nature, yielding encouraging performance in reconstructing videos, while it is likely applicable to decoding of mental images. The encoding performance of VAE is relatively higher in early visual areas than higher visual areas, but overall not as high as the performance of CNN. In summary, the findings in this paper support the Bayesian brain hypothesis, and highlight a generative strategy for decoding naturalistic and diverse visual experiences.

### Bayesian Brain and Free energy

This work is inspired by the Bayesian brain theory (Ma et al., 2006; Yuille and Kersten, 2006). This theory has been used to model the brain in various functional domains, including cognitive development (Perfors et al., 2011; Tenenbaum et al., 2011), perception (Knill and Pouget, 2004; Lee and Mumford, 2003), and action (Friston, 2010). This theory assumes that the brain learns a generative model of the world and uses it to infer the hidden cause of sensation (Friston, 2012). Under this theory, the brain runs computation similar to Bayesian inference (Knill and Richards, 1996); visual perception is viewed as a probabilistic process (Pouget et al., 2013), capable of dealing with noisy, incomplete, or ambiguous visual input (Tenenbaum et al., 2011; Yuille and Kersten, 2006).

VAE rests on a very similar notion. The encoder computes latent variables as Gaussian distributions, from which the samples are drawn by the decoder. The latent variable distribution can be used to generate not only the samples already learned, but also samples that are unknown to the model. VAE also embodies the inference strategy called “analysis by synthesis” (Yuille and Kersten, 2006). In this regard, the top-down generative component is important for optimizing the bottom-up component, to find what cause the input. Given the sensory input, the bottom-up process makes proposals of possible causes that generate the sensation (Yuille and Kersten, 2006), and then the proposals are updated by the top-down generative model with the direct comparison between sensory input and its generated “virtual version” (Friston and Kiebel, 2009). In VAE, this notion is represented by the learning rule of minimizing the reconstruction error - the difference between the input to the encoder and the output from the decoder. This learning rule allows the (bottom-up) encoder and the (top-down) decoder to be trained all together (Kingma and Welling, 2013).

VAE also contributes to the computational approximation of “free-energy”, a neuroscience concept to measure the discrepancy between how the sensation is represented by the model, and the way it actually is (Friston, 2010; Hinton and Zemel, 1994). In Bayesian inference, minimizing this discrepancy (also called “surprise”) is important for updating the model but is difficult to calculate. Therefore free-energy is proposed as the upper-bound of the sensory surprise which can be minimized coherently by minimizing free-energy (Friston, 2009). Mathematically, free-energy is expressed as the sensory surprise plus the non-negative KL divergence between the inferred causes given the input, and the true hidden causes. This formulation simplifies the inference process to an easier optimization problem by approximating the posterior distributions of visual causes (Friston, 2010). VAE bears the same idea in an artificial neural network. As shown in Eq. (1), the learning objective *L*(***θ, φ; x***) can be decomposed into two parts: the marginal likelihood of the data log*p*_***θ***_(***x***), minus the KL divergence between inferred latent variables and its ground truth *D*_*KL*_[*q*_***φ***_(***z |x***)||*p*_***θ***_(***z |x***)]. As the marginal likelihood is the negative format of “surprise” (Friston et al., 2012), VAE is actually optimizing the probability of generated data through minimizing “free-energy”.

### VAE vs. CNN

Our results suggest that VAE, as an implementation of Bayesian brain, turns out to partially explain and decode brain responses to natural videos. Therefore, it lends support to the Bayesian brain theory. However, its encoding and decoding performance is not perfect, or even worse than CNN, which outperforms VAE in nearly all visual areas. While both trained with the same data with different learning objective, the difference in encoding performance between VAE and CNN was most notable in higher visual areas in the ventral stream, which has been known to play an essential role in object/scene recognition (Ungerleider and Haxby, 1994). CNN is explicitly driven by object categorization, because its training is supervised by categorical labels of images. VAE is trained without using any label, thereby with unsupervised learning.

It is likely that supervised learning is required for a model to fully explain ventral-stream activity. A previous study has reached a similar conclusion with different models and an analysis method based on representational similarity (Khaligh-Razavi and Kriegeskorte, 2014). However, this conclusion should be taken with caution. There are many potential learning objectives that can be used for unsupervised learning. The argument on unsupervised versus supervised learning still awaits future studies to fully resolve.

### Towards Robust and Generalizable Natural Visual Reconstruction

Researchers have long been trying to render evoked brain activities into the sensory input, including edge orientation (Kamitani and Tong, 2005), face images (Cowen et al., 2014; Güçlütürk et al., 2017; Nestor et al., 2016), contrast patterns (Miyawaki et al., 2008), digits (Du et al., 2017; Qiao et al., 2018; van Gerven et al., 2010b) and handwritten characters (Schoenmakers et al., 2013). Other studies have tried to extend the reconstruction from artificial stimuli to diverse natural stimuli. The reconstruction of diverse natural images or movies has been shown by using the Bayesian reconstruction method (Naselaris et al., 2009; Nishimoto et al., 2011), where the reconstruction is matched to the images or movies in the prior dataset. Recently deep generative models have been used for decoding natural vision. A deep convolutional generative adversarial network (DCGAN) (Radford et al., 2015) enables the reconstruction of handwritten characters and natural images (Seeliger et al., 2017). Besides, a deep generator network (DGN) (Nguyen et al., 2016) has been introduced to add naturalistic constraints to image reconstruction based on decoded human fMRI data (Shen et al., 2017).

Our method is different by its training strategy. Unlike the Bayesian reconstruction method, VAE-based decoding model is generalizable without relying on the natural image prior. Besides, both DCGAN and DGN are optimized with adversarial training (Goodfellow et al., 2014). Whereas adversarial training may constrain visual reconstruction to be naturalistic (Shen et al., 2017), the VAE-based decoding model is purely rooted in variational inference. In addition, our goal is not only to optimize neural decoding for visual reconstruction, but also to test the plausibility of using VAE to model the brain. Although VAE and GAN are both emerging techniques for learning deep generative models (Dumoulin et al., 2016), these two types of networks are not mutually exclusive; instead, they might be combined for potentially better and more generalizable visual reconstruction.

### Limitations and Future Directions

Although our results support the Bayesian brain theory, some limitations of VAE should be noted. VAE offers a computational account of Bayesian inference and free-energy minimization in the brain. However, the biological inference might be implemented in a more complex and dynamic process. In this sense, the brain infers hierarchically organized sensory causes possibly with predictive coding (Huang and Rao, 2011; Rao and Ballard, 1999). Higher-level neural systems attempt to predict the inputs to lower-level ones, and prediction errors of the lower-level are propagated to adapt higher-level systems to reduce the prediction discrepancy (Clark, 2013). Therefore, hierarchical message passing with recurrent and feedback connections might be essential for effective implementation of Bayesian inference in the brain (Bastos et al., 2012; Friston, 2010; Friston and Kiebel, 2009). While VAE cannot offer hierarchically organized sensory causes and temporal processing, these properties might be important and should be incorporated into the future development of similar models.

## Acknowledgements

The authors would like to recognize the inputs from Dr. Eugenio Culurciello, Dr. Alfredo Canziani and Abhishek Chaurasia for their constructive discussions on deep neural networks. The research was supported by NIH R01MH104402 and Purdue University.

